# Genome-wide association study of glucocerebrosidase activity modifiers

**DOI:** 10.1101/2024.03.27.586821

**Authors:** Emma N. Somerville, Lynne Krohn, Konstantin Senkevich, Eric Yu, Jamil Ahmad, Farnaz Asayesh, Jennifer A. Ruskey, Dan Spiegelman, Stanley Fahn, Cheryl Waters, S. Pablo Sardi, Roy N. Alcalay, Ziv Gan-Or

**Affiliations:** The Neuro (Montréal Neurological Institute-Hospital), McGill University, Montréal, QC, Canada; Department of Human Genetics, McGill University, Montréal, QC, Canada; Department of Neurology and Neurosurgery, McGill University, Montréal, QC, Canada; Department of Neurology, College of Physicians and Surgeons, Columbia University Medical Center, New York, NY, USA; Rare and Neurological Diseases Therapeutic Area, Sanofi, Cambridge, MA, USA; Taub Institute for Research on Alzheimer’s Disease and the Aging Brain, Columbia University Medical Center, New York, NY, USA

**Keywords:** genome wide association study, GWAS, Parkinson’s disease, glucocerebrosidase, GAA, GBA1, lysosomal metabolism.

## Abstract

One of the most common genetic risk factors for Parkinson’s disease (PD) are variants in *GBA1*, which encodes the lysosomal enzyme glucocerebrosidase (GCase). GCase deficiency has been associated with an increased PD risk, but not all individuals with low GCase activity are carriers of *GBA1* mutations, suggesting other factors may be acting as modifiers. We aimed to discover common variants associated with GCase activity, as well as replicate previously reported associations, by performing a genome-wide association study using two independent cohorts: a Columbia University cohort consisting of 697 PD cases and 347 controls and the Parkinson’s Progression Markers Initiative (PPMI) cohort consisting of 357 PD cases and 163 controls. As expected, *GBA1* variants have the strongest association with decreased activity, led by p.N370S (beta = -4.36, se = 0.32, p = 5.05e-43). We also identify a novel association in the *GAA* locus (encoding for acid alpha-glucosidase, beta = -0.96, se = 0.17, p = 5.23e-09) that may be the result of an interaction between GCase and acid alpha-glucosidase based on various interaction analyses. Lastly, we show that several PD-risk loci are potentially associated with GCase activity. Further research will be needed to replicate and validate our findings and to uncover the functional connection between acid alpha-glucosidase and GCase.

## 1. Introduction

Parkinson’s disease (PD) is a complex neurodegenerative disorder, characterized by accumulation of alpha synuclein in Lewy bodies and a progressive loss of dopaminergic neurons in the substantia nigra [1]. Although the exact mechanisms remain unknown, there is a clear role for genetic factors in PD. One gene of particular interest is *GBA1*. In addition to being one of the most common genetic risk factors associated with PD, it is also associated with a faster rate of motor and non-motor progression [2].

*GBA1* encodes for glucocerebrosidase (GCase), a lysosomal hydrolase whose biallelic deficiency causes the lysosomal storage disorder (LSD) known as Gaucher disease [3]. Heterozygous *GBA1* variants associated with reduced GCase activity are important risk factors for both PD and dementia with Lewy bodies (DLB), and a substantial portion of patients with these disorders have reduced GCase activity albeit not carrying *GBA1* variants [4, 5]. This observation suggests that other factors, genetic or environmental, may modify GCase activity. One such factor may be *TMEM175* variants, which have been associated with reduced GCase activity in patient data [6] and in cell models [7]. Additionally, variants in *LRRK2* have also been linked to modified GCase activity, although the directionality of the effect is still unclear [4, 8–10].

Understanding the genetic variation influencing GCase activity could be informative in several ways. First, it can provide a better understanding of the mechanisms underlying the role GCase plays in PD. Furthermore, it would allow for proper adjustments or stratification in future research and clinical trials. In the present study, we aimed to identify genetic factors associated with GCase activity by performing a genome-wide association study (GWAS) including a total of 1054 PD cases and 510 healthy controls. A secondary analysis was conducted to identify PD-associated variants that may act as modifiers of GCase activity.

## 2. Methods

### 2.1 Study Population

The study population consisted of two separate cohorts with available genotype and GCase activity data: 1) a cohort of 697 PD cases and 347 controls collected from Columbia University in New York, and 2) a cohort of 357 PD cases and 163 controls from the Parkinson’s Progression Markers Initiative (PPMI). Cohort demographics can be found in Table 1. Both cohorts have been previously described [4, 9]. All subjects were of European descent, confirmed with principal component analysis. Informed consent forms were signed by all participants prior to entering their respective studies and the study protocol was approved by the institutional review boards.

**Table 1.** Cohort demographics for individuals with available genotype and GCase data.

| Cohort | N Cases | N Controls | Status | Males (%) | Females (%) | Mean Age (±SD) |
| --- | --- | --- | --- | --- | --- | --- |
| Columbia | 697 | 347 | cases | 65.1 | 34.9 | 65.74 (11.05) |
|  |  |  | controls | 38.3 | 61.7 | 64.25 (10.05) |
| PPMI | 357 | 163 | cases | 66.7 | 33.3 | 61.85 (9.49) |
|  |  |  | controls | 67.5 | 32.5 | 60.79 (11.34) |

N, number; SD, standard deviation; PPMI, Parkinson's Progression Markers Initiative.

### 2.2 Enzyme Activity

Enzymatic activity for GCase, acid alpha-glucosidase (GAA), acid sphingomyelinase (ASM), alpha-galactosidase A (GLA), and galactosylceramidase (GALC) was measured from dried blood spots in the Columbia cohort, the protocol for which has been previously described [11, 12]. To summarize, enzyme activity was measured by Sanofi laboratories using liquid chromatography-tandem mass spectrometry (LC-MS/MS) from dried blood spots, using a multiplex assay. The dried blood spots were incubated in a reaction cocktail with substrates for the enzymes and buffer. Enzyme activity measurements were determined by calculating the amount of product obtained after the incubation, under the assumption that this is a direct indication of the activity of the enzymes. Lysosomal enzyme activity for the PPMI cohort was measured from frozen whole blood, which was slowly thawed and the processed similarly to the Columbia cohort. GCase outliers were identified as those with activity measurements lying outside +/- 1.5 times the inter quartile range (IQR), and were subsequently removed in both cohorts.

### 2.3 Genome-Wide Association Study

Genotyping was performed with the OmniExpress GWAS array for the Columbia cohort and the NeuroX array for the PPMI cohort, according to manufacturer’s protocols (Illumina Inc.). Quality control for individual and variant data was completed as previously described (https://github.com/neurogenetics/GWAS-pipeline). In brief, samples that were heterozygosity outliers (inclusion criteria of -0.15 <= F <= 0.15), call rate outliers (missingness > 95%), had mismatched genetic and reported sex, or were identified as European ancestry outliers based on HapMap3 principal component analysis (PCA) in plink v1.9 were removed [13]. Additionally, we removed samples with relatedness closer than 3^rd^ degree relatives (pihat > 0.125). Individual SNPs were excluded on the basis of variant missingness (> 95%), differences in missingness between cases and controls (p < 1e-04), haplotype missingness (p < 1e-04), and deviation from Hardy-Weinberg equilibrium in controls (p < 1e-04). Imputation was performed on filtered data with the Michigan Imputation Server using the Haplotype Reference Consortium reference panel r1.1 2016 and default settings [14]. Linear regressions of GCase activity were performed in plink v1.9 using hard-call variants (R^2^ > 0.8) and a minor allele frequency (MAF) threshold of > 0.01. We manually added the PD-relevant variant *GBA1* p.N370S to the Columbia analyses, as well as *GBA1* p.T369M and *LRRK2* p.G2019S after filtering, due to all being under the minor allele threshold. The Columbia cohort included adjustments for age, sex, disease status, Ashkenazi Jewish ancestry, *LRRK2* p.G2019S status, and lysosomal enzyme activities (GAA, GLA, GALC, and ASM) to examine the isolated effect on GCase activity. The p.G2019S mutation was included as a covariate due it being a known GCase modifier [4, 8, 9]. Lysosomal enzymes were adjusted for because they have been shown to correlate with one another in a previous study [15]. The PPMI analysis included the same adjustments, with the addition of white blood cell count (WBC) as previously suggested [9]. Fixed-effect meta-analyses of PPMI and Columbia cohorts were performed with METAL in plink v1.9. Principal components (PCs) were calculated using PCA with plink v1.9 and the top 10 PCs were included as covariates for each cohort. Conditional and joint analyses were performed with GTCA-COJO to identify independent variants after adjusting for lead SNPs [16]. ANOVA, linear regressions with interaction term, and interaction plots were created in R v4.3.1 [17]. Linkage disequilibrium Manhattan plots were created using LocusZoom [18].

### 2.4 Data Statement

All code for analyses used in this project can be found at https://github.com/gan-orlab/GCase_GWAS. PPMI data used for this study can be obtained by qualified researchers upon completion of a data access application (https://www.ppmi-info.org/access-data-specimens/download-data). Columbia cohort data can be obtained by request. The GCase GWAS summary statistics can be found on the GWAS catalog (https://www.ebi.ac.uk/gwas/).

## 3. Results

### 3.1 *GAA* and *GBA1* loci are associated with GCase activity

After quality control and imputation, a total of 556 cases, 284 controls and 13327381 variants in the Columbia cohort and 328 cases, 144 controls and 941882 variants in the PPMI cohort were available for analysis. Mean GCase activity was similar between Columbia (mean = 11.1 umol/l/h, sd = +/- 3.14) and PPMI (mean = 11.63 umol/l/h, sd = +/- 2.59).

We performed a GWAS to identify potential associations between common variants and GCase activity in both cohorts separately, followed by a meta-analysis. We looked at the genomic inflation factors and QQ plots of each analysis to test for systematic bias, and found them to be acceptable (Columbia λ = 0.99, PPMI λ = 1.04, Supplementary Figure 1). The strongest associated locus in Columbia was in the *GBA1* locus, as expected (Figure 1A). This signal is driven by p.N370S (rs76763715, beta = -4.21, se = 0.35, p = 4.55e-31), which is in linkage disequilibrium (LD) with the lead SNP in the locus (rs745550122, beta = -4.46, se = 0.35, p = 6.61e-34, R^2^ = 0.78, D’ = 0.91). A second signal can be seen in chromosome 17 in the *GAA* locus, with the strongest associated variants comprising a high-LD region (Supplementary Figure 2) and the lead SNP having a negative direction of effect (beta = -0.96, se = 0.17, p = 7.55e-9). Applying conditional and joint analyses revealed no secondary independent associations in either signal. The *GBA1* signal was replicated in the PPMI cohort, with p.N370S once again driving the association (Figure 1B, beta = -5.05, se = 0.76, p = 1.25e-10). The *GAA* locus does surpass nominal significance in PPMI, although the lead SNP has an opposite direction of effect compared to the lead SNP from Columbia (Table 2). The meta-analysis likewise showed associations in the *GBA1* and *GAA* loci, but did not result in any additional associations (Figure 1C). The *GBA1* variants p.T369M (rs75548401) and p.E326K (rs2230288) were also associated with GCase activity, although not at GWAS significance (Table 3). Both variants had similar effect sizes, directions, and p-values across both analyses.

**Figure 1.**
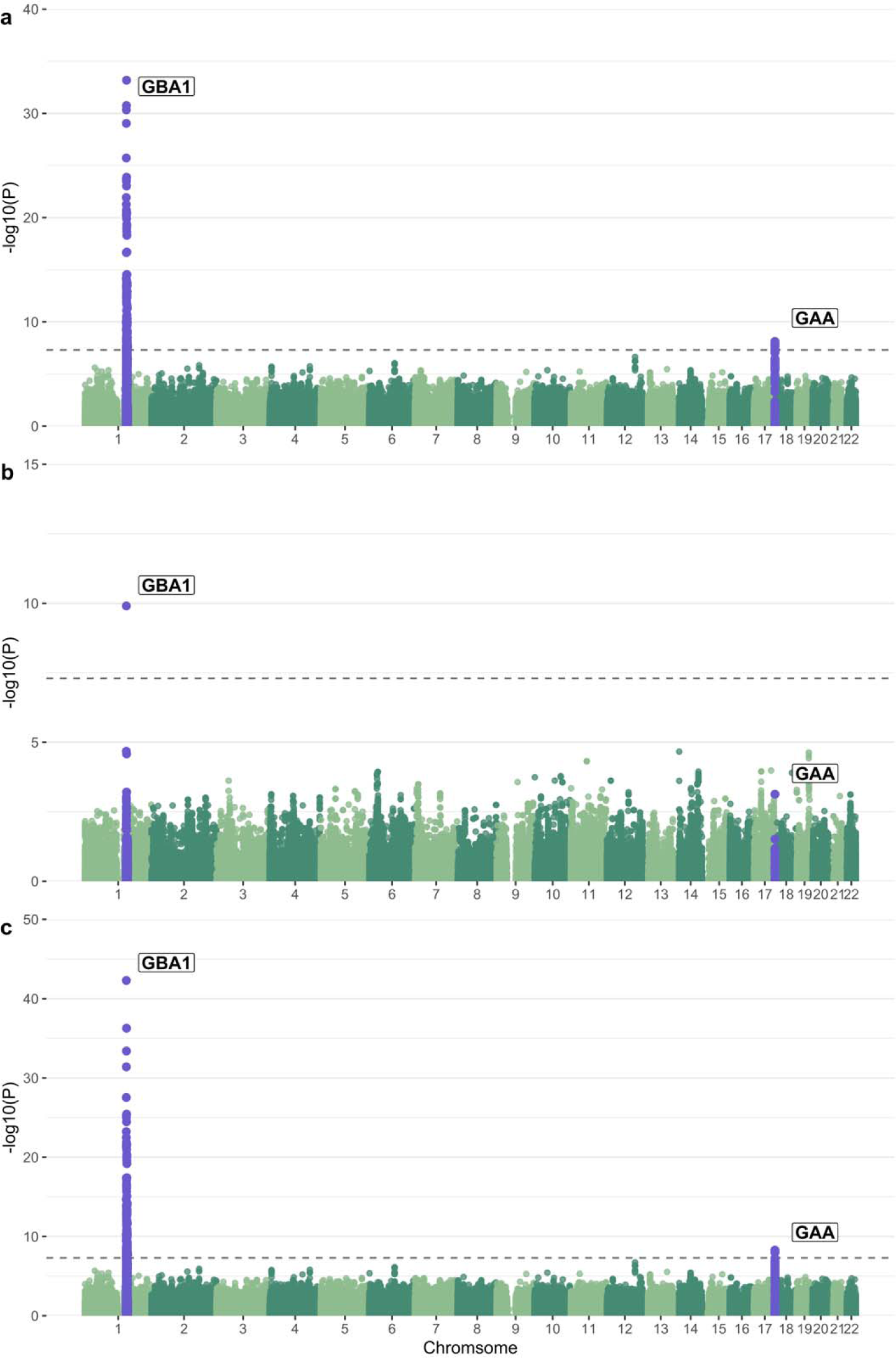
Manhattan plot of log adjusted p-values at each genomic position in a) the Columbia cohort with adjustments for age, sex, disease status, Ashkenazi Jewish status, *LRRK2* p.G2019S, ASM activity, GAA activity, GLA activity, GALC activity, and the top 10 PCs, b) the PPMI cohort with adjustments for age, sex, disease status, *LRRK2* G2019S genotype, ASM activity, GAA activity, GLA activity, GALC activity, white blood cell count, and the top 10 PCs, and c) the meta-analysis of these two analyses.

**Table 2.**
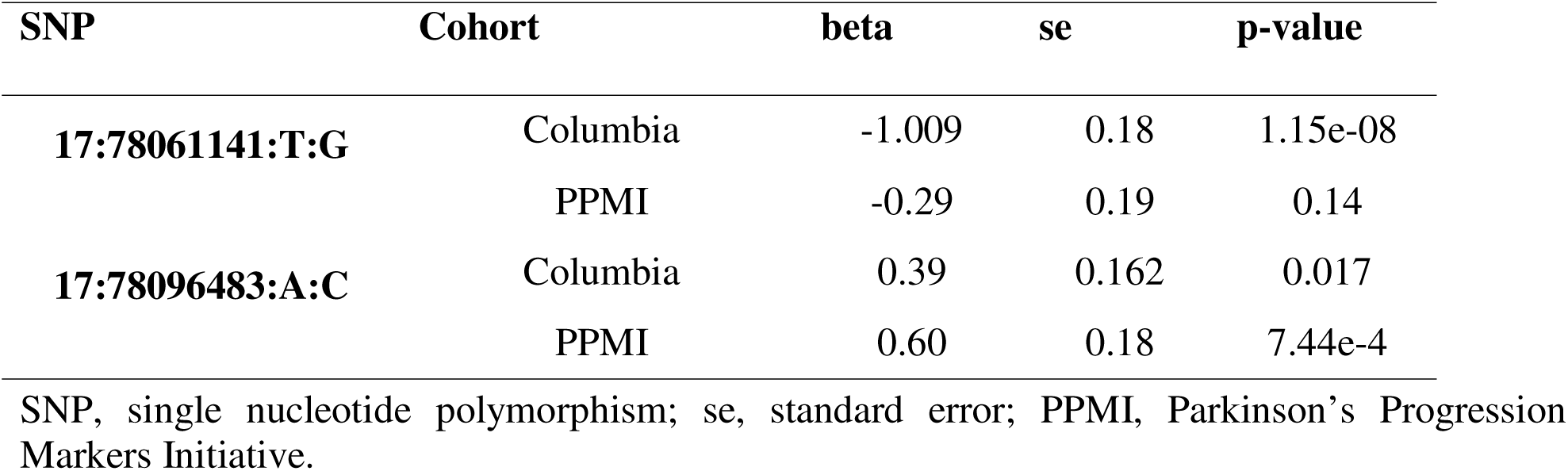
Comparison of lead *GAA* locus variants in Columbia and PPMI cohorts.

| SNP | Cohort | beta | se | p-value |
| --- | --- | --- | --- | --- |
| 17:78061141:T:G | Columbia | -1.009 | 0.18 | 1.15e-08 |
|  | PPMI | -0.29 | 0.19 | 0.14 |
| 17:78096483:A:C | Columbia | 0.39 | 0.162 | 0.017 |
|  | PPMI | 0.60 | 0.18 | 7.44e-4 |

SNP, single nucleotide polymorphism; se, standard error; PPMI, Parkinson's Progression Markers Initiative.

**Table 3.** *GBA1* p.T369M and p.E326K association statistics for Columbia and PPMI cohorts.

| Variant | Cohort | beta | se | p-value |
| --- | --- | --- | --- | --- |
| p.T369M | Columbia | -3.28 | 0.78 | 2.79e-05 |
|  | PPMI | -2.91 | 0.68 | 2.11e-05 |
|  | Meta-analysis | -3.067 | 0.51 | 1.787e-09 |
| p.E326K | Columbia | -1.62 | 0.59 | 0.0063 |
|  | PPMI | -1.43 | 0.5 | 0.0047 |
|  | Meta-analysis | -1.51 | 0.38 | 8.027e-05 |

se, standard error; PPMI, Parkinson's Progression Markers Initiative.

Interestingly, analysis in the Columbia cohort without adjusting for acid alpha-glucosidase activity eliminates any association from the *GAA* locus (Figure 2A). Adding adjustments for common PD-relevant *GBA1* variants p.N370S, p.E326K and p.T369M results in both the *GBA1* and *GAA* peaks dropping below GWAS-level significance (Figure 2B). To better understand if these outcomes could be due to an interaction between the acid alpha-glucosidase and GCase enzymes, we constructed an interaction plot of enzyme activities by genotype of the top *GAA* SNP that was common in both cohorts (Figure 2B). The genotype of the lead SNP in the *GAA* locus (17:78061141:T:G) appears to impact the interaction of GCase and acid alpha-glucosidase enzyme activities in Columbia, with correlations becoming progressively weaker in an additive manner with the addition of the T allele (Figure 3). This trend is not as clear in controls, which could be due to the lower number of samples in this group. The effect is also not apparent in PPMI, reflective of the lack of association in this cohort. To further investigate the presence of an interaction, both an ANOVA and a linear regression using an interaction term were performed. The ANOVA used the absolute difference between GCase and acid alpha-glucosidase activity, which did significantly vary by the genotype status of the lead *GAA* variant (df = 2, Sum Sq = 130, Mean Sq = 64.75, F = 13.62, p = 1.52e-06). The linear regression was performed using the same covariates as the previous analysis, with the addition of a *GAA*\*GCase interaction term. The results indicated that the T/T genotype status was significant (beta = -0.25, se = 0.13, p = 0.048), supporting the presence of an interaction effect between *GAA* genotype status, acid alpha-glucosidase activity, and GCase activity.

**Figure 2.**
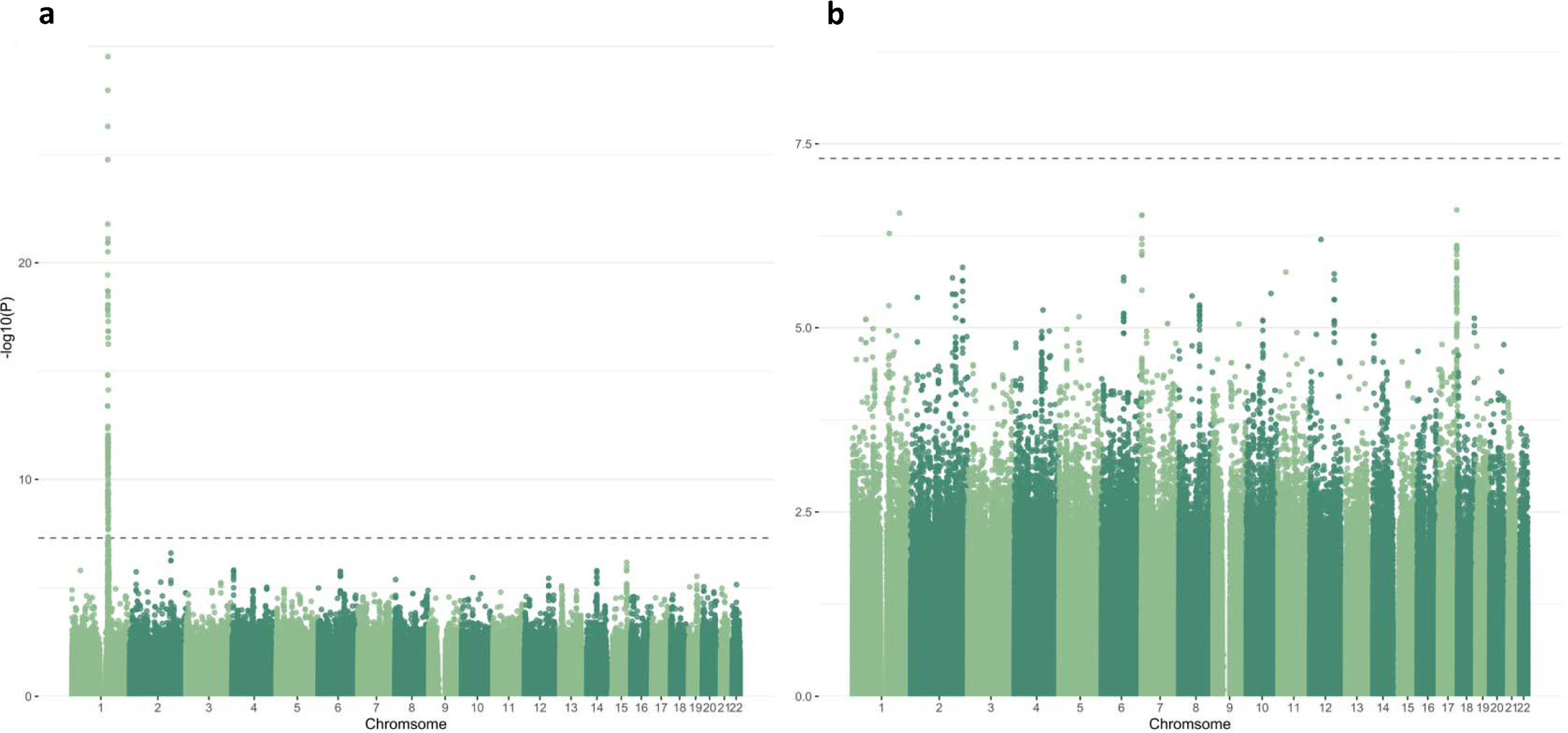
Manhattan plot of log adjusted p-values at each genomic position in the Columbia cohort with adjustments for a) age, sex, disease status, Ashkenazi Jewish status, *LRRK2* p.G2019S, ASM activity, GLA activity, GALC activity, and the top 10 PCs, and b) age, sex, disease status, Ashkenazi Jewish status, *LRRK2* p.G2019S, *GBA1* p.N370S, *GBA1* p.T369M, *GBA1* p.E326K, ASM activity, GAA activity, GLA activity, GALC activity, and the top 10 PCs.

**Figure 3.**
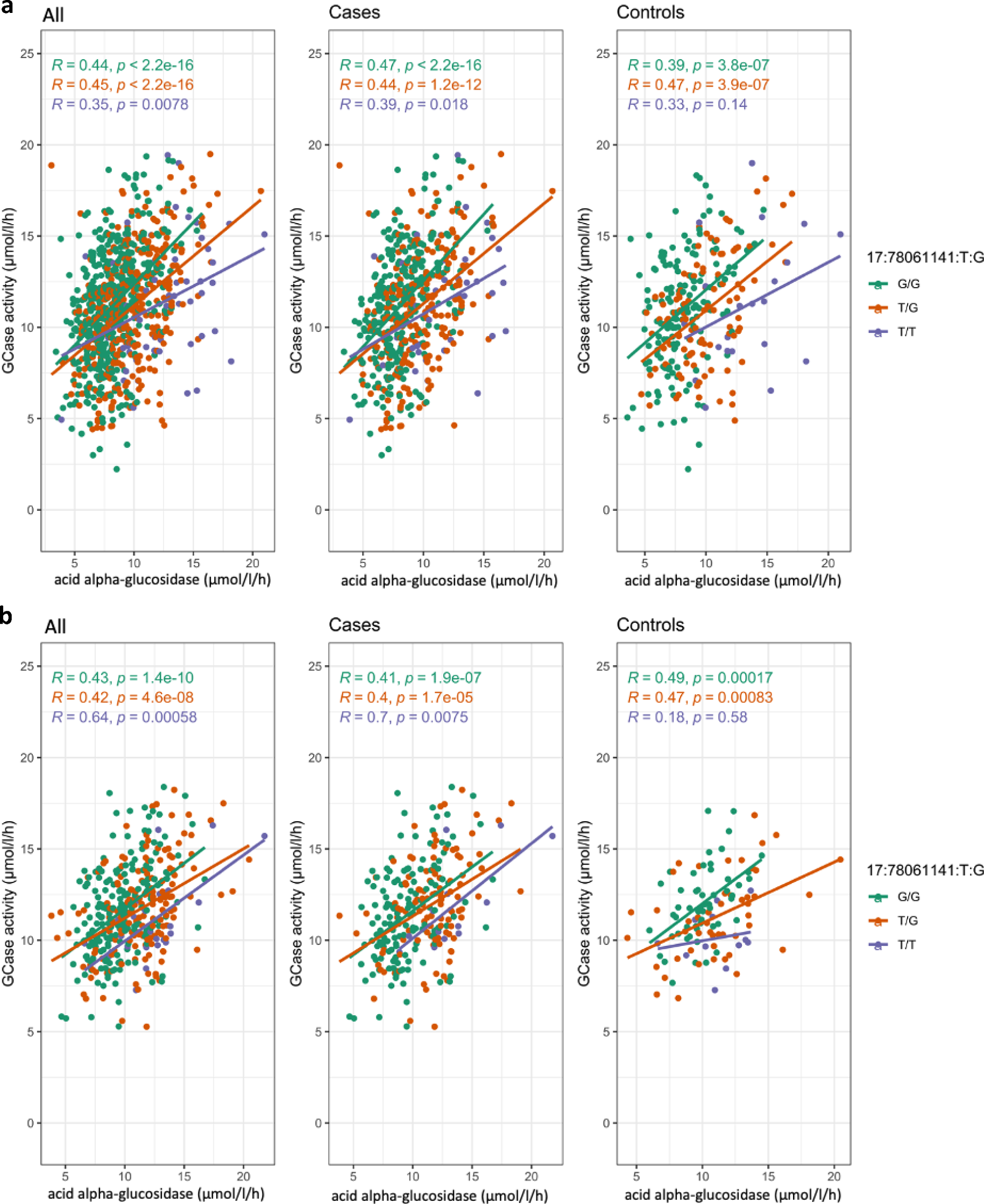
Interaction plots of GCase and acid alpha-glucosidase activity colored by 17:78061141:T:G genotype in a) Columbia and b) PPMI

To examine if the *GAA* locus has any role in PD, we investigated the association of this region with PD risk and various measurements of progression using GWAS summary statistics from previous studies. We identified 26, 4, 5, 79, and 6 variants that were nominally associated in the GWASs of PD risk, age at onset, UPDRS III, MoCA, and MMSE scores, respectively (Supplementary Table 1), although no variants passed Bonferroni correction for multiple comparisons [19–21]. There were no *GAA* locus variants even nominally associated in the GWASs of motor, cognitive, and composite progression [22]. Of the associated variants, a total of 8, 2, 33, and 1 variants in the GWASs of PD risk, age at onset, MoCA, and MMSE scores were also nominally associated with GCase activity in the present meta-analysis.

### 3.2 PD-related variants are associated with GCase activity

We investigated PD-related variants nominated from the largest PD GWAS to date to determine if any were associated with GCase activity in our cohorts (Nalls et al., 2019). There were 17, 7 and 14 loci below nominal significance in the main GWAS for Columbia, PPMI and the meta-analysis, respectively (Table 4). Of these, only p.N370S (rs76763715) and rs35749011 (in strong LD with p.E326K) in the *GBA1* locus, as well as rs13117519 in the *ANK2* locus passed multiple testing correction in the meta-analysis. We then attempted to replicate previously reported associations in *TMEM175* and *LRRK2* with GCase activity, by investigating these loci in a simplified meta-analysis using only age, sex, disease status, 10 PCs, and PD-relevant *GBA1* SNPs including p.N370S, p.E326K, and p.T369M as covariates (Supplementary Figure 3). There were 3 intronic variants in *LRRK2* that passed nominal significance (rs116911375, beta = -1.7, se = 0.79, p = 0.03; rs191242488, beta = -1.041, se = 0.52, p = 0.044; 12:40668909:T:G, beta = - 1.041, se = 0.52, p = 0.044), and no variants in *TMEM175*. No variants passed multiple testing correction. The nominally significant variants do not appear to be in LD with p.G2019S or p.M1646T. Although these variants are not associated with GCase activity in our analysis, p.G2019S and p.M1646T do show a consistent positive direction of effect with what has been demonstrated in previous research (Table 5) [4, 9, 10]. The difference seen in our results compared to the nominal association of p.M1646T found in a very similar analysis in these cohorts performed by Sosero et al. [10] is due to the removal of GCase outliers in our study. With the inclusion of outliers, we also see an association of this variant in our meta-analysis (b = 1.16, se = 0.38, p = 0.002, rs35303786).

**Table 4.** Parkinson’s disease risk SNPs associated with changes in GCase activity.

| Cohort | SNP | rsID | Beta | SE | MAF | P | Locus |
| --- | --- | --- | --- | --- | --- | --- | --- |
| Columbia | 1:155205634:T:C | rs76763715 | -4.21 | 0.35 | 0.041 | 4.55E-31 | *GBA1* |
|  | 20:6006041:C:T | rs77351827 | 0.63 | 0.19 | 0.16 | 0.00083 | *CRSL1* |
|  | 6:112243291:A:G | rs997368 | -0.49 | 0.16 | 0.24 | 0.0017 | *FYN* |
|  | 4:114369065:C:T | rs13117519 | 0.53 | 0.19 | 0.16 | 0.006 | *ANK2* |
|  | 4:90666041:C:T | rs356203 | 0.37 | 0.14 | 0.41 | 0.008 | *SNCA* |
|  | 4:90626111:G:A | rs356182 | 0.36 | 0.14 | 0.37 | 0.011 | *SNCA* |
|  | 1:155135036:G:A | rs114138760 | -1.61 | 0.64 | 0.012 | 0.012 | *GBA1* |
|  | 21:38852361:G:A | rs2248244 | 0.39 | 0.16 | 0.26 | 0.012 | *DYRK1A* |
|  | 4:90636630:G:A | rs5019538 | 0.33 | 0.14 | 0.33 | 0.022 | *SNCA* |
|  | 17:43933307:T:C | rs7221167 | -0.3 | 0.14 | 0.4 | 0.029 | *MAPT* |
|  | 14:37989270:T:C | rs12147950 | -0.3 | 0.14 | 0.43 | 0.032 | *MIPOL1* |
|  | 4:90607126:C:G | rs356228 | 0.28 | 0.13 | 0.46 | 0.034 | *SNCA* |
|  | 18:31304318:G:T | rs1941685 | -0.29 | 0.14 | 0.47 | 0.037 | *ASXL3* |
|  | 3:28705690:T:C | rs6808178 | 0.3 | 0.14 | 0.36 | 0.039 | *LINC00693* |
|  | 13:97865021:T:C | rs4771268 | -0.32 | 0.16 | 0.28 | 0.044 | *MBNL2* |
|  | 14:95194760:G:C | rs4905237 | 0.38 | 0.19 | 0.15 | 0.047 | *SERPINA13P* |
| PPMI | 1:155205634:T:C | rs76763715 | -5.048 | 0.76 | 0.0087 | 1.25E-10 | *GBA1* |
|  | 1:155135036:G:A | rs35749011 | -1.43 | 0.5 | 0.02 | 0.0047 | *GBA1* |
|  | 1:154898185:G:C | rs114138760 | -1.61 | 0.62 | 0.014 | 0.0097 | *GBA1* |
|  | 4:114369065:C:T | rs13117519 | 0.48 | 0.19 | 0.19 | 0.01 | *ANK2* |
|  | 1:171719769:C:T | rs11578699 | 0.47 | 0.19 | 0.19 | 0.012 | *VAMP4* |
|  | 14:88464264:G:T | rs979812 | -0.48 | 0.2 | 0.45 | 0.018 | *GALC* |
|  | 6:32578772:C:A | rs504594 | -0.46 | 0.22 | 0.14 | 0.035 | *HLA-DRB1* |
| Meta | 1:155205634:T:C | rs76763715 | -4.36 | 0.32 | 0.036 | 5.05E-43 | *GBA1* |
|  | 1:155135036:G:A | rs35749011 | -1.50 | 0.39 | 0.017 | 0.00014 | *GBA1* |
|  | 4:114369065:C:T | rs13117519 | 0.50 | 0.13 | 0.17 | 0.00016 | *ANK2* |
|  | 20:6006041:C:T | rs77351827 | 0.45 | 0.14 | 0.15 | 0.0013 | *CRSL1* |
|  | 4:90666041:C:T | rs356203 | 0.31 | 0.1 | 0.41 | 0.0028 | *SNCA* |
|  | 4:90636630:G:A | rs5019538 | 0.3 | 0.11 | 0.33 | 0.0041 | *SNCA* |
|  | 6:112243291:A:G | rs997368 | -0.34 | 0.12 | 0.22 | 0.0053 | *FYN* |
|  | 14:88464264:G:T | rs979812 | -0.34 | 0.13 | 0.44 | 0.0068 | *GALC* |
|  | 4:90626111:G:A | rs356182 | 0.2784 | 0.1037 | 0.3811 | 0.007286 | *SNCA* |
|  | 1:154898185:G:C | rs114138760 | -1.606 | 0.618 | 0.0138 | 0.009358 | *GBA1* |
|  | 21:38852361:G:A | rs2248244 | 0.3915 | 0.1556 | 0.2634 | 0.01187 | *DYRK1A* |
|  | 18:31304318:G:T | rs1941685 | -0.2872 | 0.1376 | 0.4749 | 0.03687 | *ASXL3* |
|  | 4:90607126:C:G | rs356228 | 0.2054 | 0.1005 | 0.4607 | 0.04097 | *SNCA* |
|  | 2:135438789:G:A | rs4954162 | 0.2489 | 0.1246 | 0.2193 | 0.04585 | *TMEM163* |
|  | 14:95194760:G:C | rs4905237 | 0.3794 | 0.1904 | 0.1481 | 0.0463 | *SERPINA13P* |

SNP, single nucleotide polymorphism; se, standard error; MAF, minor allele frequency; PPMI, Parkinson's Progression Markers Initiative.

**Table 5.** *LRRK2* p.G2019S and p.M1646T association statistics for Columbia and PPMI cohorts.

| **Variant** | **Cohort** | **beta** | **se** | **p-value** |
| --- | --- | --- | --- | --- |
| **p.G2019S** | Columbia | 0.64 | 0.4 | 0.11 |
|  | PPMI | -0.053 | 1.12 | 0.96 |
|  | Meta-analysis | 0.56 | 0.56 | 0.14 |
| **p.M1646T** | Columbia | 0.41 | 0.45 | 0.36 |
|  | PPMI | 0.82 | 0.54 | 0.13 |
|  | Meta-analysis | 0.58 | 0.58 | 0.094 |

se, standard error; PPMI, Parkinson's Progression Markers Initiative.

## 4. Discussion

In the present GWAS, we identified variants in the *GAA* locus as potential modifiers of GCase activity, in addition to the known associations with *GBA1* variants. *GAA* encodes acid alpha-glucosidase, and deficiency in this enzyme leads to the LSD Pompe disease [23]. There is no known association between Pompe disease and PD, and the variants in the *GAA* locus are not associated with PD risk, age at onset, or various measures of progression. This is an important observation, since it may suggest that changes in GCase activity alone are insufficient to cause PD, or that the effects of *GAA* variants on GCase activity are too small to have a meaningful clinical effect. It was hypothesized that the mechanism underlying *GBA1*-associated PD is an imbalance, or disturbance in the flux within the lysosomal glycosphingolipid metabolism pathway, rather than reduced GCase activity on its own [24]. This is supported by the identification of other genes in this pathway that are associated with PD, such as *SMPD1*, *GALC*, *ARSA* and *ASAH1* [12, 25–32]. Previous work has also demonstrated that lysosomal enzymes are highly correlated, especially for GCase, acid alpha-glucosidase, and alpha galactosidase A which is encoded by *GLA* [15]. Taken into consideration with the interaction effect observed in the present study, there seems to be mounting evidence in favor of this hypothesis. The mechanism behind the interaction of GCase and acid alpha-glucosidase is still unknown. The products of GCase are lipids and glucose, while acid alpha-glucosidase hydrolyzes glycogen into glucose [33]. Our observation of genotype-dependent correlation between GCase and acid alpha-glucosidase activities may therefore indicate that the *GAA* locus variants associated with GCase activity could maintain equilibrium in this pathway, as they affect both GCase and alpha-glucosidase activities. This hypothesis requires additional genetic, enzymatic, and functional studies.

We further examined the association between known PD-associated loci and GCase activity. As to be expected, we found three PD variants in the *GBA1* locus to be associated with GCase activity. We also found an additional association with the *ANK2* locus. Phosphorylation in the *ANK2* region has been proposed to play a role in PD neurodegeneration through the inhibition of organelle autophagy, but its connection to GCase activity is not clear from the current literature and will need further research [34]. There are also multiple notable associations with ties to GCase activity that may have only failed multiple testing correction due to sample size. One such locus is *GALC*, encoding the enzyme galactosylceramidase which, similarly to GCase, breaks down large sphingolipids into lipids and ceramide. The association of *GALC* variants with GCase activity may suggest that a disturbance in lysosomal metabolism could be a causal mechanism in PD, which is also supported by previous research which found galactosylceramide and GCase to be correlated [15]. We also found nominal associations with multiple *SNCA* variants, complimenting previous associations found between levels of alpha synuclein aggregates and Gcase activity [35–37]. A third interesting locus that fell just shy of the multiple testing correction threshold was *FYN*. This gene encodes a kinase that has been shown to phosphorylate alpha-synuclein, which is a key step towards its aggregation into Lewy bodies [38]. We did not replicate previously reported associations with *LRRK2* or *TMEM175* variants, which is likely due to additional adjustments in our study and the removal of GCase outliers compared to previous studies. Studies in larger cohorts will be required to confirm the associations of these PD loci.

This study has several limitations. The first limitation is a lack of power. Our discovery and replication analyses utilized the largest cohorts with enzyme activity measurements that we could find at the time of performing the study and allowed us to complete the first GWAS of GCase activity, but only enabled us to discover common variants with moderate effect sizes or rare variants with large effect sizes. Due to this, there are likely more modulators of GCase activity yet to be uncovered. A second limitation is the use of only individuals of European and Ashkenazi Jewish ancestry, as there was insufficient data to carry out the study in other populations. Next, our study had minor differences in age and notable differences in sex, particularly in the Columbia cohort, between cases and controls. Both age and sex were adjusted for in all analyses to account for these differences. An additional limitation is the method of enzyme measurement used. Lysosomal activity was measured using dried blood spots, which is not guaranteed to capture the true enzyme activity within a functioning lysosomal environment. These results will strengthen if replicated in future studies that use lysosomal-specific methods of enzyme activity, in cells more representative of disease pathogenesis.

In conclusion, we found a novel potential association in the *GAA* locus associated with GCase activity, which may represent an interaction effect between GCase and acid alpha-glucosidase and could be indicative of a homeostatic relationship between lysosomal enzymes. We also support previously suggested connections of multiple PD-related genes with GCase activity, such as *GALC* and *SNCA*, as well as identifying a novel potential association with *ANK2*. These findings could be significant for improving our understanding of how GCase deficiency is related to overall lysosomal dysfunction, and how GCase functions in relation to other lysosomal enzymes. Additionally, associated loci can be taken into account in future research and clinical trials to help control for genetic influences on GCase activity. Due to the novelty of our results and lack of confident association in our replication cohort, further genetic and functional research will be required to validate our findings.

## 6. Acknowledgements

This study was financially supported through grants from the Michael J. Fox Foundation (MJFF), the Canadian Consortium on Neurodegeneration in Aging (CCNA), and the Canada First Research Excellence Fund (CFREF), awarded to McGill University for the Healthy Brains for Healthy Lives initiative (HBHL). Data used in the preparation of this article were obtained from the Parkinson’s Progression Markers Initiative (PPMI) database (www.ppmi-info.org/access-data-specimens/download-data). For up-to-date information on the study, visit www.ppmi-info.org. PPMI – a public-private partnership – is funded by the Michael J. Fox Foundation for Parkinson’s Research and funding partners, including [list the full names of all of the PPMI funding partners found on the PPMI Website]. We would like to thank the research participants for contributing to this study. We thank Meron Teferra for her assistance. Z.G.O. is supported by the Fonds de recherche du Québec—Santé (FRQS) Chercheurs-boursiers award and is a William Dawson Scholar. KS is supported by a post-doctoral fellowship from the Canada First Research Excellence Fund (CFREF), awarded to McGill University for the Healthy Brains for Healthy Lives initiative (HBHL). The Columbia cohort was funded by the National Institutes of Health (K02NS080915, and UL1 TR000040, formerly the National Center for Research Resources, Grant Number UL1 RR024156).

