## Supplementary Figures for "Genome-wide association study of glucocerebrosidase activity modifiers"

**Supplementary Figures**
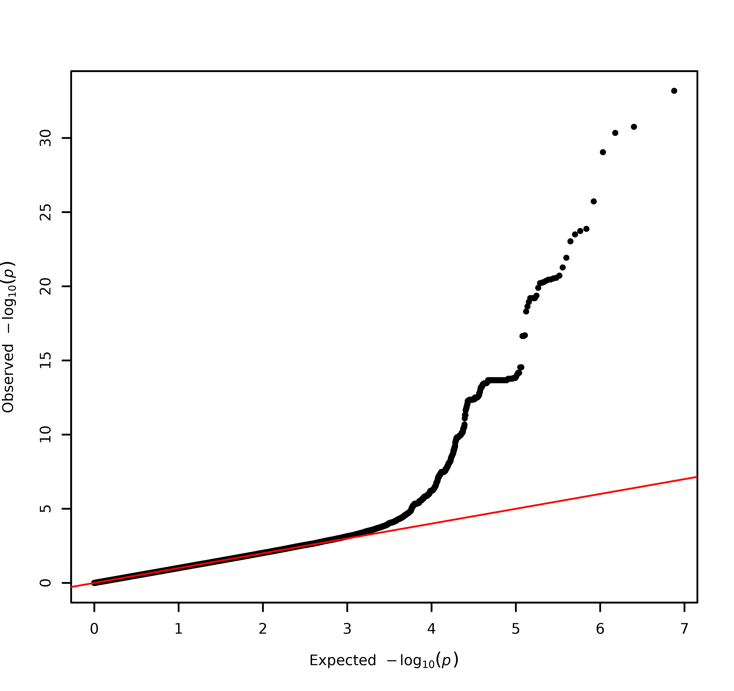

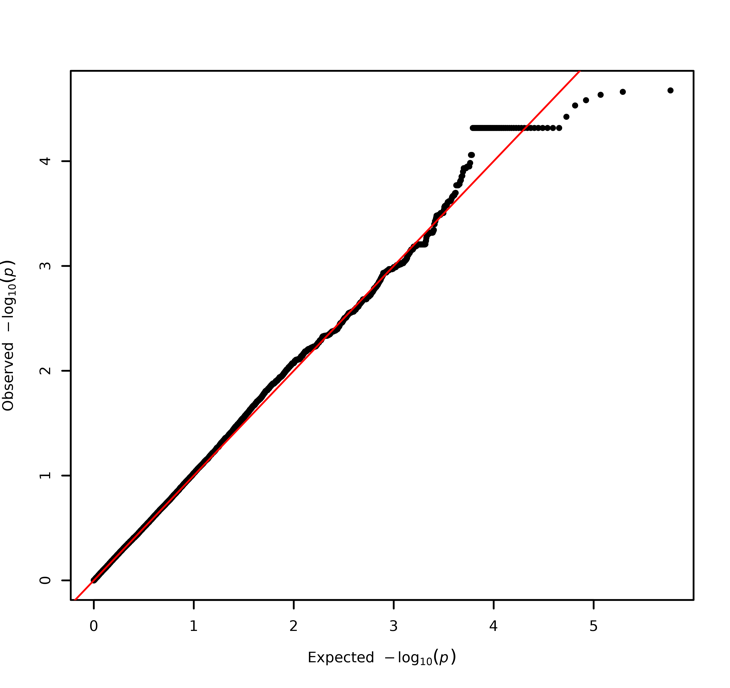

**a**

**b**

**Supplementary Figure 1.** QQ plots of linear regression p-values for a) the Columbia cohort with adjustments for age, sex, disease status, Ashkenazi Jewish status, *LRRK2* p.G2019S, ASM activity, GAA activity, GLA activity, GALC activity, and the top 10 PCs, b) the PPMI cohort with adjustments for age, sex, disease status, *LRRK2* G2019S genotype, ASM activity, GAA activity, GLA activity, GALC activity, white blood cell count, and the top 10 PCs

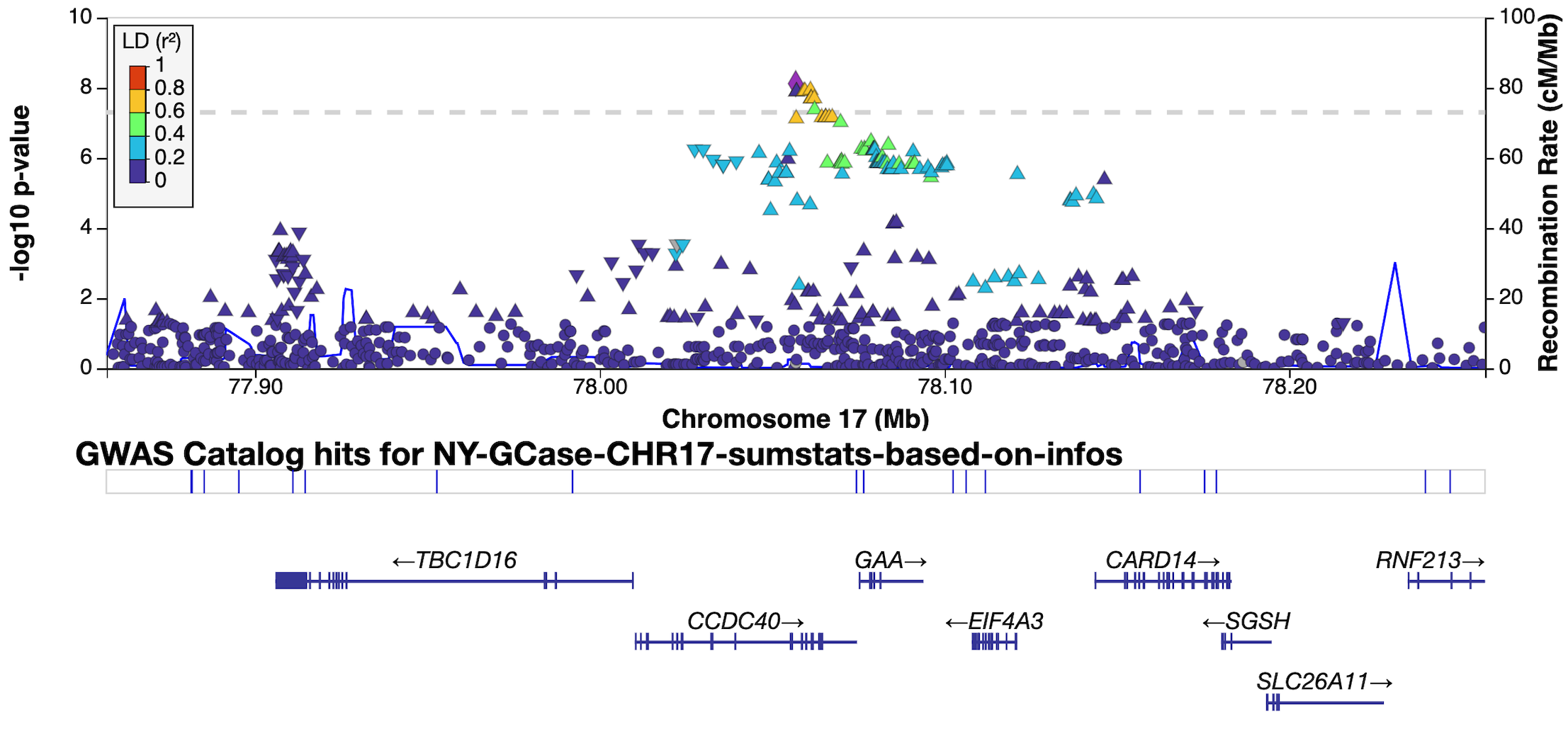

**Supplementary Figure 2**. LocusZoom Manhattan plot of the *GAA* locus from the analysis of the Columbia cohort. Color is described in the legend and based on linkage disequilibrium (R^2^).

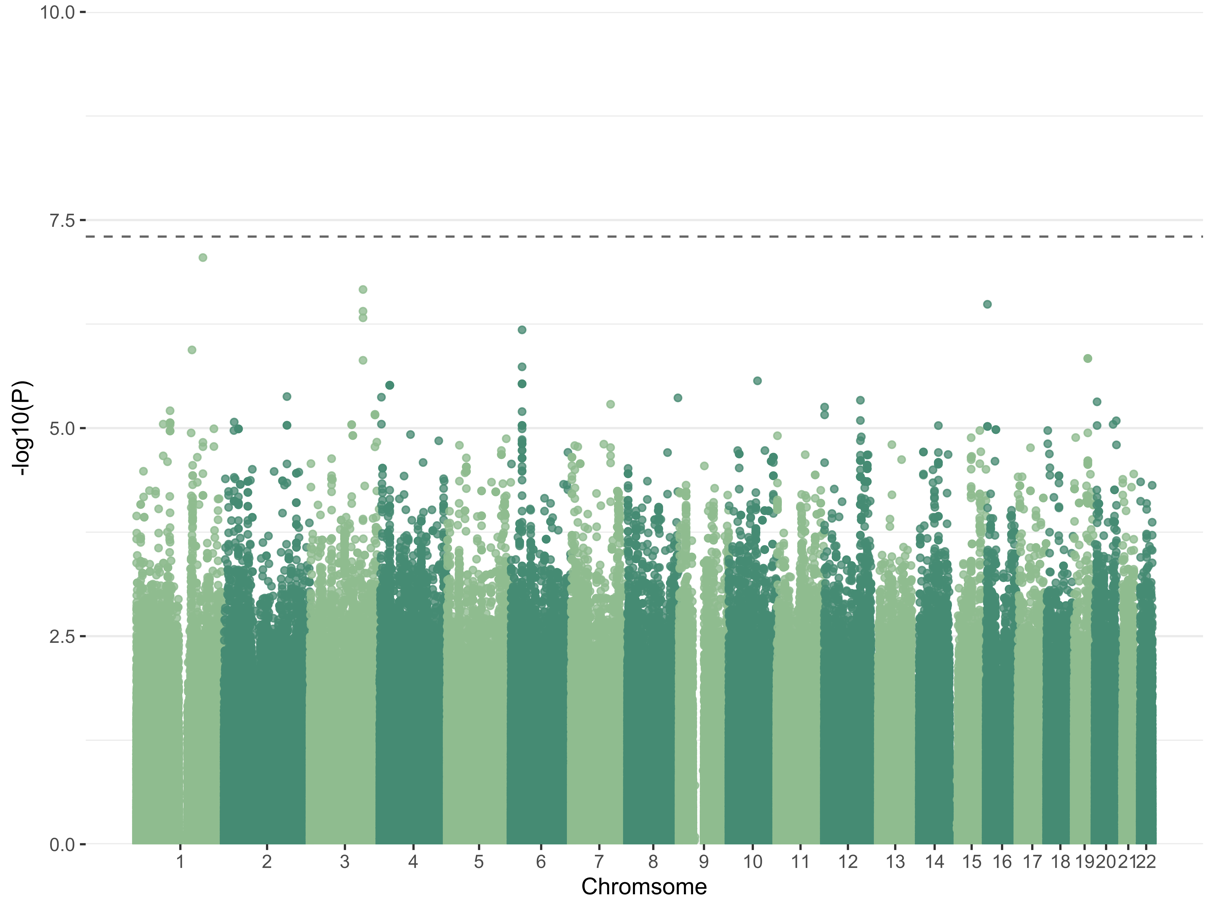

**Supplementary Figure 3**. Meta-analysis of Columbia and PPMI cohorts using age, sex, disease status, *GBA1* p.N370S, p.E326K, and p.T369M genotype status, and 10 PCs.
